# A multimodal AI system for out-of-distribution generalization of seizure detection

**DOI:** 10.1101/2021.07.02.450974

**Authors:** Yikai Yang, Nhan Duy Truong, Jason K. Eshraghian, Christina Maher, Armin Nikpour, Omid Kavehei

## Abstract

Epilepsy is one of the most common severe neurological disorders worldwide. The International League Against Epilepsy (ILAE) define epilepsy as a brain disorder that generates (1) two unprovoked seizures more than 24 hrs apart, or (2) one unprovoked seizure with at least 60% risk of recurrence over the next ten years. Complete remission has been defined as ten years seizure free with the last five years medication free. This requires a cost-effective ambulatory ultra-long term out-patient monitoring solution. The common practice of self-reporting is inaccurate. Applying artificial intelligence (AI) to scalp electroencephalogram (EEG) interpretation is becoming increasingly common, but other data modalities such as electrocardiograms (ECGs) are simpler to collect and often recorded simultaneously with EEG. Both recordings contain biomarkers in the detection of seizures.

Here, we propose a state-of-the-art performing AI system that combines EEG and ECG for seizure detection, tested on clinical data with early evidence demonstrating generalization across hospitals. The model was trained and validated on the publicly available Temple University Hospital (TUH) dataset. To evaluate performance in a clinical setting, we conducted nonpatient-specific inference-only tests on three out-of-distribution datasets, including EPILEPSIAE (30 patients) and the Royal Prince Alfred Hospital (RPAH) in Sydney, Australia (31 patients shortlisted by neurologists and 30 randomly selected). Across all datasets, our multimodal approach improves the area under the receiver operating characteristic curve (AUC-ROC) by an average margin of 6.71% and 14.42% for prior state-of-the-art approaches using EEG and ECG alone, respectively. Our model’s state-of-the-art performance and robustness to out-of-distribution datasets can improve the accuracy and efficiency of epilepsy diagnoses.

## I. Introduction

Epilepsy affects about 1% of people globally, placing it as one of the most common severe neurological disorders worldwide [1]. It entails some severe neuropsychiatric and psycho-social comorbidities that include stigmatization, social exclusion, and unemployment. The side-effects of daily medicine consumption include depression and anxiety [2]–[5]. Accurate and objective seizure counting play an integral role in a wide range of clinical diagnoses and management decisions for epilepsy.

The International League Against Epilepsy (ILAE) define epilepsy as a brain disorder that generates (1) two unprovoked seizures that are more than 24 hrs apart, or (2) one unprovoked seizure with at least 60% risk of recurrence over the next ten years [6]. Accurate seizure counting in the long-term has important implications for driving the management of epilepsy [6]. For instance, being seizure-free for ten years, while off antiepileptic drugs (AEDs) for at least five, identifies whether epilepsy is considered in remission. Epilepsy misdiagnosis or delayed diagnosis is unfortunately still common and has serious consequences [7], [8]. False positives can lead to inappropriate prescription of AEDs that result in mistreated or worsening symptoms [8], [9]. This issue is compounded by societal inequities, as 80% of patients with epilepsy are amongst low to middle-income populations, and 75% of them do not receive any treatment [10]. The treatment gap can be attributed to inequities in distribution and access to services, stigma associated with the disease, lack of sufficient expert resources (neurologists), and an inadequate supply of AEDs.

The golden standard of epilepsy diagnosis relies on surface or scalp electroencephalogram (EEG) readings and accurate annotation [11]. Over the past two decades, there has been widespread use of EEG signals for seizure documentation and seizure forecasting [12], [13]. Plausible reasons for the abnormally high level of false positives of clinician-readings may include the lack of formal standards or mandatory training in reading EEG, the time consuming nature of analyzing lengthy recordings, and the absence of confirmatory readings. A strong case can be made for secondary EEG interpretations given the difficulties outlined, and is an opportunity for automated seizure detection models to act as clinician support systems. Beyond augmented intelligence systems, fully autonomous agents performing seizure detection is a prerequisite to developing closed-loop neurostimulation devices for seizure suppression.

Recently, several deep learning techniques have achieved promising results using EEG for non-patient-specific seizure detection on the Temple University Hospital (TUH) EEG dataset [14]. Beyond clinical use (in- and out-patients), EEG-based methods are limited as the recording apparatus is typically not designed for ongoing wear and would otherwise cause discomfort. Attempts to reduce the number of EEG channels have yielded limited results. A recent approach saw the Neureka 2020 Epilepsy Challenge accounting for the number of channels in their scoring formula. Despite this, the winner of this challenge relied on a 16-channel EEG and still only managed to achieve 12.37% sensitivity (with one false alarm per 24 hours) [15]. 16-channels are clearly inappropriate for ambient use, and the state-of-the-art result highlights the challenges associated with developing a high-performance EEG-based seizure detection system using a constrained number of electrodes. Portability requires complementary biomarkers to EEGs that are already integrated into wearable devices [16].

Epileptic seizures are known to cause short-term and longterm heart rate disturbances. Ictal tachycardia occurs in over 80% of partial onset-seizures [17], [18], which may precede electrographic or clinical onset [19], [20]. Compared with EEG, ECGs are relatively more portable, and are routinely recorded simultaneously with EEG [21]. Most successful studies using ECG have focused on identification of seizure onset (early prediction) [22], [23], for instance, using heart-rate variability (HRV) to predict events in children with temporal-lobe epilepsy [19], [24]–[26]. In contrast, ECG for seizure detection has had limited focus as its performance is not yet comparable with multi-channel EEGs. Despite the lower performance of lone usage, ECG recordings can also extract markers of seizure events that EEGs may not pick up.

On clinical utility, a significant challenge in AI-based seizure detection is the need for models that generalize across patients, recording equipment, and hospitals. The reality is that many clinics are unlikely to generate a sufficient amount of labeled data in a format immediately usable as a training set for deep learning models. The practices of one clinic are likely to differ from those of another, as is the recording equipment and parameters (e.g., variation in sampling rates, slight differences in electrode positioning). There is no guarantee that the large epilepsy datasets available can generalize sufficiently enough to deploy hospital-specific models. Unfortunately, all prior works reported in a recent review [27] on the application of deep learning on seizure detection, are retrospective and were only benchmarked on test sets sourced identically to the training set. Such models provide low confidence for deployment in clinics that differ from where the data was gathered. This requires rigorous testing and high performance on out-of-distribution datasets. Despite being a key barrier to deployment, generalization across hospitals is not a common metric that is optimized for due to its associated difficulties.

The recent study in [28] leveraged the abundance of weak annotations that were analyzed by a mixed group of technicians, fellows, students, and epileptologists to train a convolutional neural network (CNN), achieving an area under the receiver operating characteristic curve (AUC-ROC) score of 0.78. When generalizing the network to the Stanford hospital dataset, the AUC-ROC score dropped to 0.70. This study provides one of very few publicly available inter-hospital results of a deep learning algorithm using EEG recordings. In our recent work [29], our approach reached an AUC-ROC score of 0.84 using a convolutional long short-term memory (ConvLSTM) network tested on unseen patient data. These results show the potential of attaining specialist-level diagnostic capability that can be used as either a primary diagnostic tool or secondary decision support system. But not only is there insufficient evidence of useful out-of-distribution performance, these deep learning methods ignore insight provided by other biomarkers that clinicians have access to.

This study aims to improve the seizure detection rate in adults by combining simultaneously acquired EEG and ECG recordings in a fused deep learning system. Our study demonstrates that using both recordings in an appropriately structured multimodal neural network can provide a more robust diagnosis than either measurement alone, and also improves upon previously reported state-of-the-art multimodal neural networks applied to this task [30]. To the best of our knowledge, only a set of limited works concatenates EEG and ECG recordings for seizure detection, one in particular, achieving an AUC-ROC improvement of 0.01 and 0.11, when compared with EEG-only and ECG-only models, respectively [31], [32]. However, naively concatenating different features in a machine learning model poses several challenges. The use of more features opens up susceptibility to overfitting; combining heterogeneous sources of data increases the difficulty of feature extraction; the inability to isolate the noise of correlated distributions can increase the bias of the network. These issues can severely harm the out-of-distribution performance on unseen patients and datasets.

In this paper, we demonstrate state-of-the-art performance of non-patient-specific seizure detection, and provide early evidence of generalization on out-of-distribution datasets across continents with different data acquisition hardware infrastructure. We achieve this by designing a multimodal neural network model that accounts for the independent contributions of EEG and ECG towards the classification of seizure events, in addition to their correlated contribution. We expect this novel network architecture to be capable of improving other multimodal tasks as well. The main contributions of this paper are summarized below:

- A non-patient specific multimodal deep learning model using both EEG and ECG data is developed. We train and validate the model on the publicly available TUH dataset (US). The model is evaluated on a carved out test set, achieving an improvement of the AUC-ROC by a margin of 2.18% over the use of EEGs alone, and by 1.67% over previously state-of-the-art multimodal neural networks that use both EEG and ECG signals together.
- Early evidence of generalization is shown by pseudo-prospectively assessing performance on datasets acquired across hospitals in two different continents without any retraining. The European EPILEPSIAE dataset (30 patients) and the Australian RPAH datasets (total of 61 patients in two separate groups) are acquired with different electrophysiological recording infrastructure. The AUC-ROC on the European dataset is 0.8595, and 0.8549 for the Australian dataset. The average improvement of the AUC-ROC across all three out-of-distribution datasets over the prior state-of-the-art deep learning using EEG and ECG are 6.71% and 14.42%, respectively.
- We introduce a novel and robust multimodal deep learning model which advances upon the previous state-of-the-art in modality fusion, ‘EmbraceNet’ [30]. Our model utilizes both independent and cross-modal relationships between the two input data distributions. We develop a separate baseline using EmbraceNet and maintain a higher AUC-ROC by a margin of 1.76%.
- We evaluate the performance of our network when either EEG or ECG data modalities are missing in order to assess the usefulness of our model for patients that lack either measurement. Our proposed model performance drops by an average of only 1.53% when compared to the dedicated ECG-only baseline, and improves by an average of 1.30% when compared to the EEG-only baseline. These values are obtained by averaging the AUC-ROC of the TUH dataset and the out-of-distribution datasets, showing that our network is still effective in the absence of either recording.

To the best of our knowledge, this is the first inference-only (pseudo-prospective) study combining EEG and ECG modalities for seizure detection using a fusion of two deep learning networks. At the time of writing, this approach achieves the highest AUC-ROC on the TUH dataset. The remainder of the paper is organized as follows. The following section describes the features of the datasets used in the models. Section III introduces the proposed method for automatic seizure detection and the multimodal network. Finally, we discuss the results and conclude the paper.

## II. Dataset

Three datasets are used in this work: the Temple University Hospital (TUH) seizure corpus [14], EPILEPSIAE [33], and two sets from the Royal Prince Alfred Hospital (RPAH) [29]. All datasets contain surface EEG data from adult individuals living with epilepsy. These datasets are summarized in Tables I, II, and III. Only the TUH dataset is used to train and validate our network. Pseudo-prospective evaluations are performed on the EPILEPSIAE and RPAH datasets (without any re-training). The TUH (US), EPILEPSIAE (Germany), and RPAH (Australia) datasets are from three continents recorded with different data acquisition infrastructures.

**TABLE I:** TUH dataset summary (EEG and ECG)

| Attribute | Train/Dev (80/20) | Test |
| --- | --- | --- |
| Files | 4141 | 953 |
| Sessions | 1095 | 228 |
| Patients | 540 | 46 |
| Files with seizures | 746 | 258 |
| Sessions with seizures | 301 | 94 |
| Patients with seizures | 174 | 36 |
| Number of seizures | 2129 | 650 |
| Background duration (hours) | 669.3 | 149.2 |
| Seizure duration (hours) | 43.0 | 14.6 |
| Total duration (hours) | 712.3 | 163.8 |

\*The default training set was randomly split with a ratio of 80-20 to create a validation subset. The dedicated development set from the TUH seizure corpus was used as the carved-out test set.

**TABLE II:** The EPILEPSIAE scalp-EEG (ECG) dataset.

| Patient | Gender | Age | SN | Seizure-Foci | Surgery | RD (h) | Mean SD (s) | Range SD (s) |
| --- | --- | --- | --- | --- | --- | --- | --- | --- |
| 1 | M | 36 | 11 | Central&Parietal | N | 164.7 | 78.3 | [54.5, 162.0] |
| 2 | F | 46 | 8 | Temporal | N | 177.4 | 52.9 | [4.3, 71.3] |
| 3 | M | 41 | 8 | Temporal | Y | 143.3 | 58.6 | [34.3, 102.8] |
| 4 | F | 67 | 5 | Temporal | N | 167.8 | 121.0 | [84.3, 166.1] |
| 5 | F | 52 | 8 | Frontal&Temporal | N | 266.4 | 92.3 | [11.3, 379.1] |
| 6 | M | 65 | 8 | Temporal | N | 135.4 | 50.2 | [0.0, 105.1] |
| 7 | M | 36 | 5 | Temporal | Y | 118.1 | 46.7 | [29.3, 79.6] |
| 8 | M | 26 | 22 | Frontal | N | 115.6 | 20.1 | [0.9, 96.3] |
| 9 | M | 47 | 6 | Temporal | Y | 94.0 | 71.4 | [65.9, 83.8] |
| 10 | M | 44 | 11 | Frontal&Temporal | Y | 138.0 | 39.9 | [0.0, 68.1] |
| 11 | M | 48 | 14 | Central&Temporal | N | 138.1 | 63.6 | [19.4, 100.4] |
| 12 | M | 28 | 9 | Temporal | Y | 159.7 | 41.4 | [31.2, 60.8] |
| 13 | M | 46 | 8 | Frontal&Temporal | N | 158.1 | 91.0 | [55.0, 125.5] |
| 14 | F | 62 | 6 | Temporal | N | 162.2 | 124.0 | [83.0, 174.1] |
| 15 | F | 41 | 5 | Temporal | N | 118.7 | 64.9 | [7.6, 144.0] |
| 16 | F | 15 | 6 | Temporal | Y | 92.9 | 56.1 | [2.5, 99.0] |
| 17 | F | 17 | 9 | Temporal | N | 159.1 | 55.8 | [33.3, 76.5] |
| 18 | M | 47 | 7 | Temporal | N | 178.2 | 41.7 | [22.7, 69.4] |
| 19 | M | 32 | 22 | Temporal | N | 161.1 | 65.2 | [43.1, 96.5] |
| 20 | M | 47 | 7 | Temporal | Y | 164.6 | 59.5 | [19.1, 119.8] |
| 21 | F | 31 | 8 | Occipital | Y | 159.4 | 51.9 | [9.3, 118.9] |
| 22 | M | 38 | 7 | Parietal | Y | 137.9 | 95.7 | [55.5, 145.1] |
| 23 | M | 50 | 9 | Temporal | N | 237.5 | 56.1 | [18.8, 88.0] |
| 24 | F | 54 | 10 | Temporal | Y | 94.4 | 72.9 | [49.6, 122.8] |
| 25 | M | 42 | 8 | Central | N | 159.7 | 329.3 | [73.8, 909.2] |
| 26 | M | 13 | 9 | Temporal | Y | 159.0 | 62.7 | [38.0, 104.5] |
| 27 | M | 58 | 9 | Temporal | Y | 159.5 | 95.3 | [43.9, 331.2] |
| 28 | F | 35 | 9 | Temporal&Parietal | N | 162.3 | 86.2 | [59.4, 216.9] |
| 29 | M | 50 | 10 | Temporal | N | 161.1 | 180.7 | [62.1, 270.0] |
| 30 | F | 16 | 12 | Temporal | N | 159.8 | 75.2 | [10.0, 151.9] |
| Total | — | — | 238 | — | — | 4604 | 75.9 | [0.0, 909.2] |

M: Male, F: Female, SN: Number of seizures, RD: Recording duration, Mean SD: Mean of seizure duration, Range SD: Seizure duration range

**TABLE III:** Summary of the RPAH neurologist-selected dataset

| Patient | Gender | Age | SN | Seizure-Foci | Severity of motor activity | RD (h) | Mean SD (s) | Range SD (s) |
| --- | --- | --- | --- | --- | --- | --- | --- | --- |
| 1 | M | 20 | 10 | Generalised | Moderate | 86.9 | 13 | [7.2, 17.6] |
| 2 | M | 33 | 4 | Generalised | Minor | 93.6 | 91.9 | [51.2, 117.1] |
| 3 | M | 22 | 9 | Generalised | Moderate | 8.0 | 76.4 | [26.2, 191.0] |
| 4 | F | 41 | 1 | Generalised | Severe | 57.8 | 47.5 | [47.5, 47.5] |
| 5 | M | 21 | 7 | Generalised | Moderate | 71.9 | 423.1 | [21.9, 1324.4] |
| 6 | F | 22 | 6 | Frontal | Moderate | 97.7 | 74.5 | [57.1, 95.3] |
| 7 | M | 39 | 11 | Frontal | Moderate | 47.8 | 72.5 | [32.1, 132.3] |
| 8 | M | 38 | 2 | Frontal | Minor | 16.4 | 43.5 | [40.2, 46.7] |
| 9 | M | 62 | 7 | Frontal | Severe | 73.2 | 98.4 | [40.1, 158.6] |
| 10 | M | 31 | 7 | Frontal | Moderate | 34.7 | 63 | [17.4, 102.0] |
| 11 | F | 51 | 11 | Frontotemporal | Severe | 93.4 | 161.2 | [92.5, 175.4] |
| 12 | M | 25 | 8 | Frontotemporal | Moderate | 93.0 | 77.5 | [42.4, 154.6] |
| 13 | F | 43 | 8 | Frontotemporal | Minor | 90.1 | 178.7 | [28.0, 1095.9] |
| 14 | F | 22 | 5 | Frontotemporal | Minor | 138.2 | 79.4 | [62.5, 105.2] |
| 15 | M | 31 | 11 | Frontotemporal | Severe | 82.4 | 73.1 | [50.8, 202.7] |
| 16 | F | 58 | 14 | Frontotemporal | Moderate | 68.3 | 102.2 | [17.0, 237.7] |
| 17 | M | 41 | 8 | Temporal | Minor | 97.0 | 44 | [29.9, 69.6] |
| 18 | M | 39 | 9 | Temporal | Minor | 91.3 | 67.5 | [45.0, 78.1] |
| 19 | M | 51 | 7 | Temporal | Minor | 162.4 | 69.9 | [13.9, 147.5] |
| 20 | F | 41 | 14 | Temporal | Minor | 90.2 | 126.5 | [45.1, 719.7] |
| 21 | F | 32 | 9 | Temporal | Minor | 45.8 | 74.7 | [10.9, 118.1] |
| 22 | M | 32 | 8 | Temporal | Minor | 92.2 | 67.3 | [41.9, 135.0] |
| 23 | M | 33 | 8 | Temporal | Minor | 87.4 | 59.8 | [41.3, 113.1] |
| 24 | M | 32 | 9 | Temporal | Minor | 64.4 | 71.4 | [17.4, 138.3] |
| 25 | M | 41 | 8 | Temporal | Minor | 72.4 | 57.1 | [31.4, 112.7] |
| 26 | F | 25 | 11 | Parietal | Moderate | 92.5 | 134.7 | [10.5, 1074.6] |
| 27 | F | 48 | 2 | Parietal | Minor | 91.8 | 401.9 | [320.7, 483.1] |
| 28 | M | 28 | 4 | Parietal | Minor | 65.8 | 213.4 | [58.3, 485.8] |
| 29 | F | 24 | 6 | Focal vertex | Moderate | 113.0 | 24.8 | [3.5, 95.7] |
| 30 | M | 21 | 3 | Left hemisphere focal | Moderate | 87.9 | 98.7 | [88.7, 107.7] |
| 31 | F | 45 | 11 | Left insula | Minor | 88.2 | 38.3 | [9.7, 46.4] |
| Total | — | — | 238 | — | — | 2495.7 | 97.2 | [3.5, 1324.4] |

M: Male, F: Female, SN: Number of seizures, RD: Recording duration, Mean SD: Mean of seizure duration, Range SD: Seizure duration range

For a rigorous evaluation on challenging data, such as a variety of seizure types, and various foci on the brain network inputs to the autonomic nervous system [34], the RPAH dataset is divided into two test sets: one set includes 31 patients with different seizure foci and are shortlisted (selected) by neurologists from a long list of patients. The aim was to consider seizure type and semiology in expert selection of patients. The second set includes 30 patients who are randomly selected from the same pool without any prior information.

### A. TUH dataset

The world’s largest open database, the Temple University Hospital (TUH) seizure corpus [14], was used for training and validation of our deep learning model. The TUH dataset consists of simultaneously recorded EEG and ECG data. Patients with missing ECG recordings were omitted, leaving 1,095 sessions with 540 patients (174 participants with seizures) in the training set. The training set was randomly split (80/20) for training and validation. After training and parameter tuning, the model is then fixed for all future evaluation, hence a pseudo-prospective analysis can be undertaken.

For testing on the TUH dataset, we used 228 sessions with 46 patients (36 patients with seizures). In the absence of a publicly released labeled test set, we treat the TUH development dataset as the unseen test set to reduce bias in our assessment. This is summarized in Table I. To assess clinical utility, strictly no further training, tuning, or model selection took place beyond what we were satisfied with on the TUH train and development sets. This strictly inference-only approach on the out-of-distribution test sets is adopted to emulate a prospective study.

### B. EPILEPSIAE dataset

EPILEPSIAE is the largest epilepsy EEG database in Europe, containing EEG and ECG data from 275 patients [33]. In this work, we analyze scalp-EEG and ECG with 19 electrodes of 30 patients with 238 seizures and 4,604 recording hours in total. The sampling rate of the EEG is 256 Hz.

### C. RPAH dataset

We have extracted 192 adult in-patient EEG monitoring data between 2011 to 2019 at RPA Hospital in Sydney, and long-listed 111 patients with seizures recorded. In this study, RPAH neurologists assist in shortlisting 31 epilepsy adults with different seizure foci. Specifically, neurologists were asked to select patients with the six most common seizure types, namely generalized, frontal, frontotemporal, temporal, parietal, and unspecified focal epilepsy (see Table III). The total number of seizures and the mean seizure duration are 238 and 97.2 seconds, respectively. To confirm the reliability of our fused network, a randomized test is also performed where 30 patients from 111 adult patients with seizures are randomly selected without any prior information. Note that the ratio of the total seizure duration (time) over the total background data for RPAH data is significantly higher than the curated TUH dataset; hence it creates a highly realistic inference-only evaluation for false positives. This is mainly due to more network exposure to noise and artifacts.

## III. Methods

### A. Pre-processing

#### 1) EEG

Prior to being passed into the neural network, eye artifacts are removed, and frequency information is extracted by using independent component analysis (ICA) [35] and short-time Fourier Transform (STFT).

The EEG signals are first split into 12-second segments, and then the ICA algorithm is applied to decompose the signal into several statistically independent components. Blind source separation (BSS) [36] is used in the ICA [35] algorithm to obtain several statistically independent topographic maps. Eq. 1 shows the working principle of BSS, where 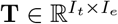 is the multi-channel EEG signal, *I_t_* represents the number of samples over time, and *I_e_* is the number of electrodes. After decomposition, 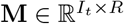 contains temporal information of the decomposed signal, 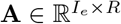 contains the topographic weight map, and *R* is the source number estimation.

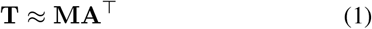

Eye movement information is recorded on the electrooculography (EOG) channel, which is physically close to the EEG channels labeled ‘FP1’ and ‘FP2’. To remove this artifact, a fully automated approach based on Pearson correlation is used [37]. Independent signal information with a strong correlation with channels ‘FP1’ and ‘FP2’ above a given threshold (based on adaptive z-scoring) are removed. STFT is then applied to the clean EEG signals with a 250 sample (or 1 second) window length and 50% overlapping. The DC component of the transform is removed as it is known to have no relation to seizure occurrences. The pre-processed dimensionality of the input data becomes (*n* × 23 × 125), where *n* is the number of electrodes, 23 is the time sample, and 125 is the range of frequencies. Artifact removal is performed using the MNE Python package. [38].

#### 2) ECG

Heart rate variability (HRV) [39] is one of the most common features extracted from ECG for seizure detection. However, for deep learning, feature engineering using HRV is unsuitable as it potentially eliminates useful information. As the ECG will ultimately be processed by a CNN, we expect the network can inherently extract relevant information without manual feature engineering. Although raw ECG signals can be directly fed into the neural network, the lack of explicit frequency information makes it difficult for the network to extract essential features. As with the EEG recordings, we apply an STFT to the ECG to translate 12 second segments of raw ECG signals into spectra as input to the neural network. To address differences in sample rates of recording equipment, all ECG signals are re-sampled to 250 Hz. Therefore, a 12 s ECG signal contains a total of 3,000 samples. We used a window length of 250 samples (or 1 s) with 50% overlapping when applying the STFT to transform the data dimensions to (23 × 1 × 126). The DC component in the spectrogram is removed, resulting in the dimensionality of (23 × 1 × 125).

### B. Deep learning network

The overall model structure can be separated into three parts. A ConvLSTM network dedicated to the EEG data, a residual CNN for the ECG data, and a fused network that takes the outputs of the individual networks to model the cross-modal representation of the EEG and ECG signals. All three networks are then connected (using both sequential and residual connections) to a terminal network consisting of several dense layers. Further details are provided in subsequent sections.

#### 1) EEG-ConvLSTM network

The deep learning network used for training the EEG signal is adopted from our previous work [29]. Three deep convolutional long short term memory (ConvLSTM) blocks [40] are combined with three fully-connected layers. The detailed structure is shown in the supplementary information Fig 3. The first ConvLSTM layer uses 16 (*n* × 2 × 3) kernels with (1 × 2) stride, where *n* represents the number of channels. The next two ConvLSTM blocks both use (1 × 2) stride and (1 × 3) kernel sizes. 32 filters and 64 for the ConvLSTM block 2 and 3, respectively. The three ConvLSTM blocks are two fully connected layers with sigmoid activation and output sizes of 256 and 2, respectively.

#### 2) ECG-ResConv network

Recently, a deep network based on convolutional neural network (CNN)-residual [41] blocks achieved excellent performance on cardiovascular disease classification problems using 12-lead ECG channels [42]. Our model fine-tunes this ResConv (CNN-residual) network to efficiently and accurately use ECG signals for seizure detection. As shown in the supplementary information Fig. 4, the input was first fed into a batch normalization layer, ensuring the input data has zero mean and unit variance to reduce the internal covariate shift [43]. The ReLU activation function was used inside the network [44], and the kernel size for all blocks was (3 × 1). The residual block was designed with a skip connection combined with two branches, and the down-sampling value in the max-pooling layer was selected to normalize the output sample sizes. The output feature size was halved block-by-block, from 64 to 8, while the number of filters was doubled block-by-block, from 32 to 256. The four residual blocks were flattened, followed by a fully connected layer with sigmoid activation and output dimension of 2. Both the flattened layer and fully connected layer had a 0.5 dropout rate.

**Fig. 1:**
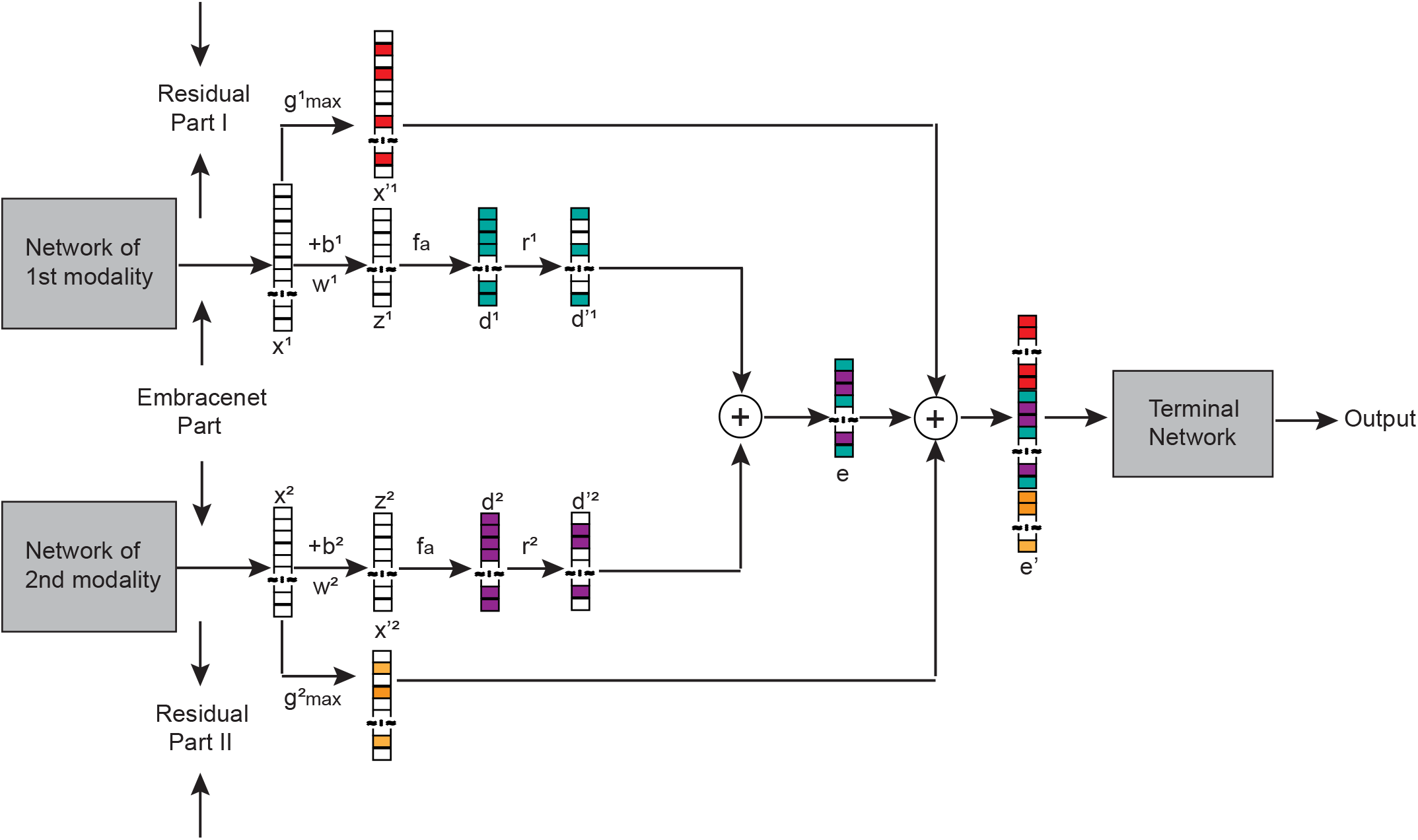
Proposed multimodal network.

**Fig. 2:**
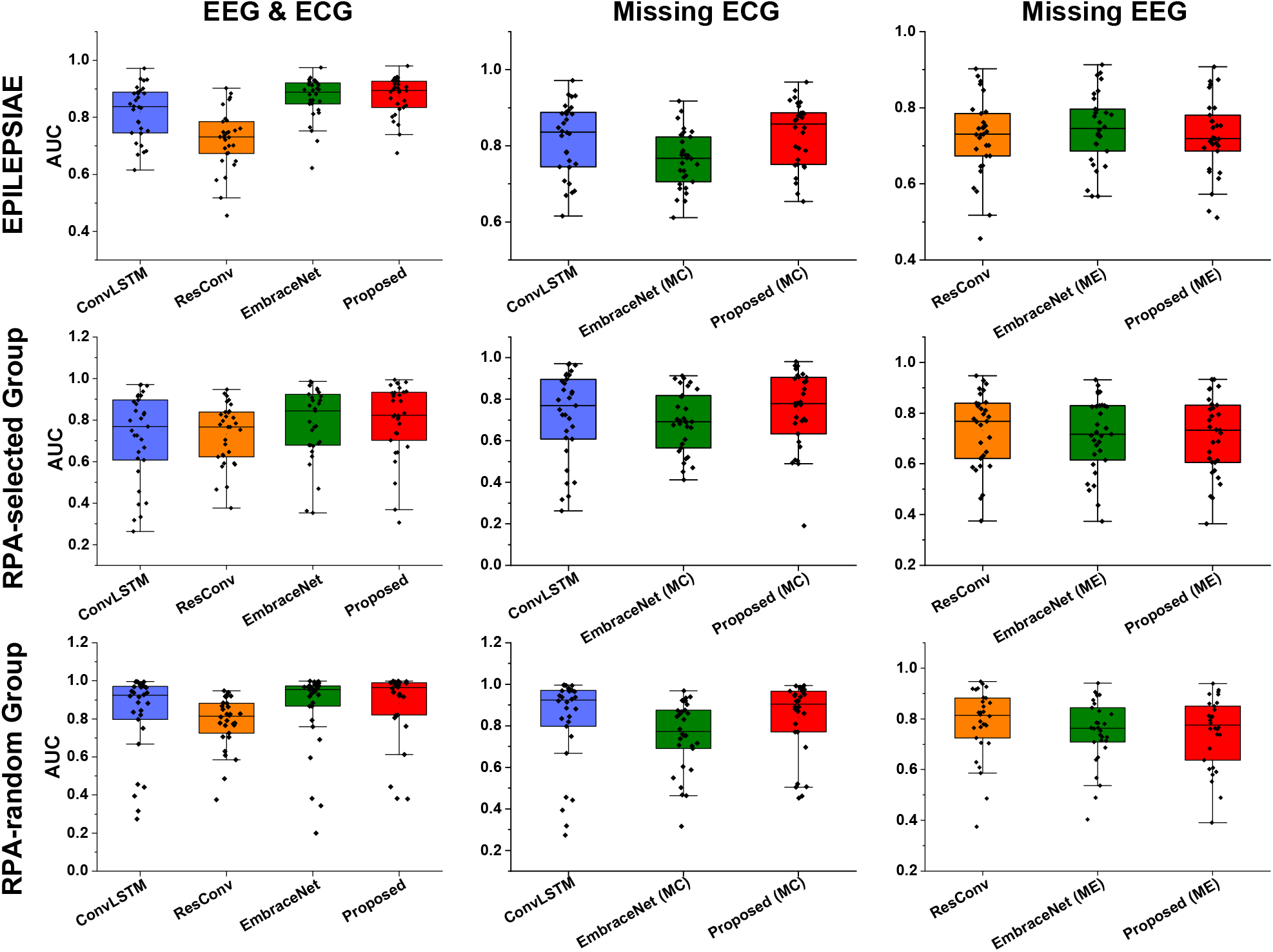
Pseudo-prospective seizure detection performance comparison. **ConvLSTM**: EEG-only with ConvLSTM network, **ResConv**: ECG-only with ResConv network, **EmbraceNet**: EmbraceNet architecture trained on combined EEG and ECG data, **Proposed**: our approach to combined EEG and ECG data, **EmbraceNet (MC)**: EmbraceNet architecture trained on EEG and ECG data, but missing ECG during inference, **EmbraceNet (ME)**: as above but missing EEG during inference, **Proposed (MC)**: our proposed approach trained on EEG and ECG data, but missing ECG data during inference, **Proposed (ME)**: as above, but missing EEG data during inference.

**Fig. 4:**
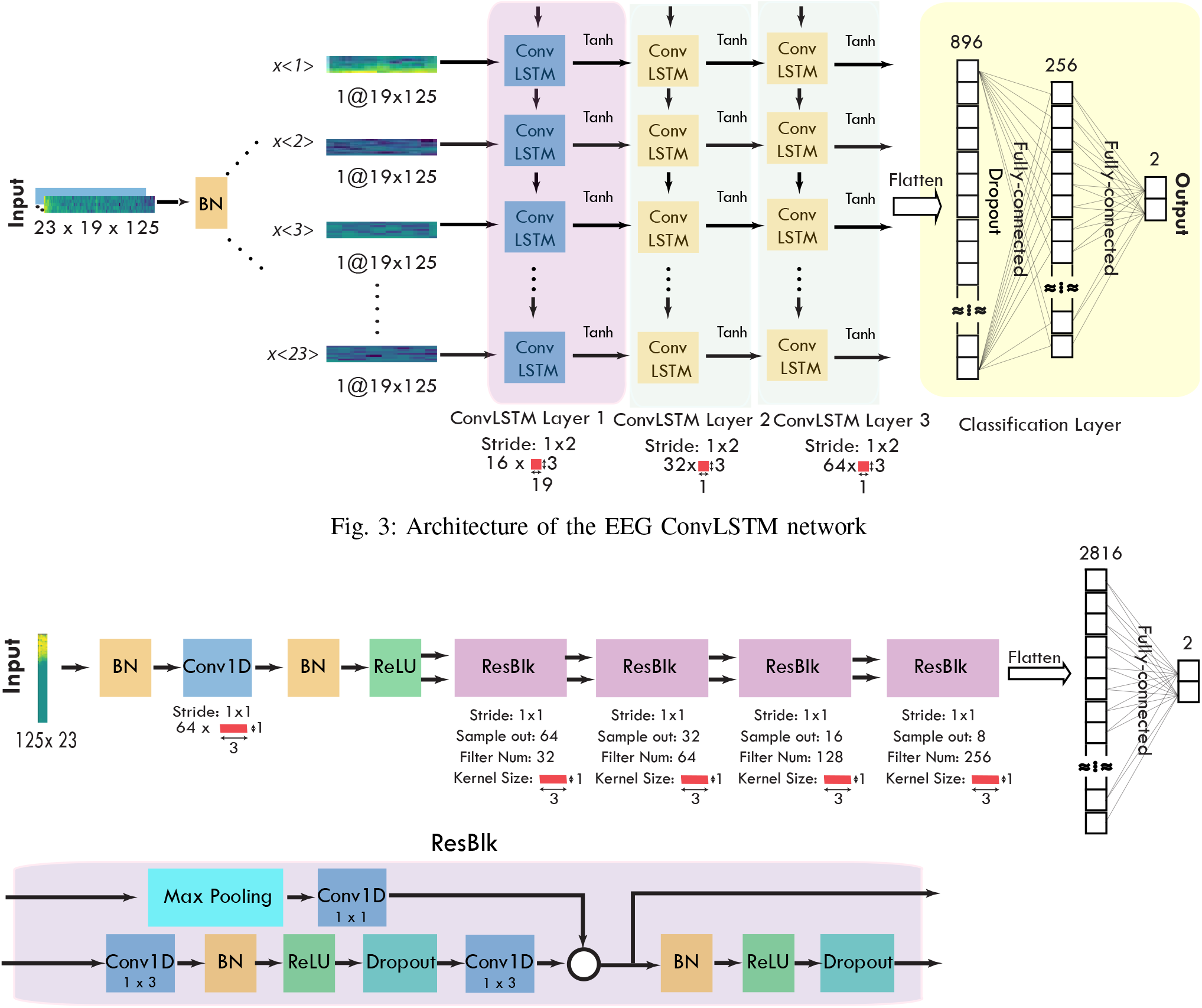
Architecture of the ECG ResConv network.

To avoid over-fitting to the training data, dropout was used and early-stopping was applied to terminate the training process when the combined training and validation loss had not decreased after 20 epochs of patience. We implemented our model in Python with Keras 2.0 with a Tensorflow 1.4.0 backend.

### C. Proposed multimodal network

#### 1) Additive representation of multimodal data distributions

The proposed network is derived from the calculation of additive signal power from a pair of correlated random variables. Multimodal data obtained from the same target is typically expected to be correlated. As an example, the average power of two superimposed waveforms containing noise content is calculated using the following equation:

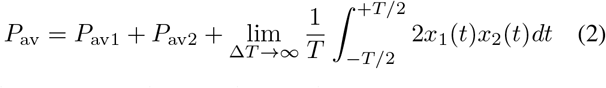

where *P*_av1_ and *P*_av2_ denote the average power of timevarying input signals *x*_1_(*t*) and *x*_2_(*t*) respectively. The third term is the correlation between the two input signals. The EmbraceNet architecture from [30] models the cross-modal correlation from the integral term, lessening the dependence on the independent contributions of the input signals present in the first two terms of Eq. 2.

To address the lack of contribution from the independent input data distributions, the EmbraceNet architecture is stacked with residual connections from the EEG-ConvLSTM network and the ECG-ResConv network (see Fig 1). This provides a more faithful representation of correlated signals, and also addresses the vanishing gradient problem in deep networks by introducing skipped pathways for gradient backpropagation.

#### 2) Network structure

The fused network takes flattened vectors *x*^(*k*)^ of the independent network models as inputs. In our case, the flattened layers of the EEG and ECG network are denoted as *x*^(1)^ and *x*^(2)^, respectively. The *i*-th input for both networks are 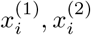.

##### a) Cross-modal correlation using EmbraceNet

The cross-modal network that finds a joint representation of the two data modalities originates from the state-of-the-art network “EmbraceNet” [30]. The outputs of the independent EEG and ECG networks are first connected to a pair of dense layers to standardize the input feature vector (*c* = 256 in our tests), reflected in the equation below:

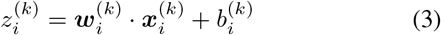

where *k* = 1, indicates the EEG network output, and *k* = 2 is the ECG network modality output. 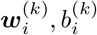 are the weights and biases, respectively.

A nonlinear activation function *f_a_* (e.g. ReLU) is applied to 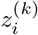, obtaining:

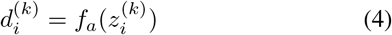

Note that *d*^(*k*)^ is of dimension *c*.

Rather than using summation to fuse the vectors, the EmbraceNet model employs an elaborate fusion technique based on a multinomial sampling process: 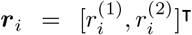 which is drawn from a multinomial distribution.

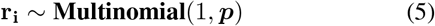

where *p* = [*p*_1_,*p*_2_]^T^ and ∑_*j*_ *P_j_* = 1, indicating that only one element of the vector *r_i_* is equal to 1, while the rest are 0. The Hadamard product between *r^k^* and *d^k^* is taken to obtain the output 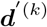 in EmbraceNet.

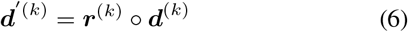

Finally, the output vector is the sum across all elements of 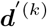.

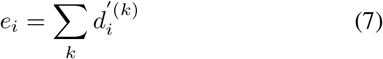

##### b) Residual Connections

To include the independent contributions of the input data distributions, residual connections are applied to the outputs of the EEG-ConvLSTM and ECG-ResConv networks (i.e., the inputs to EmbraceNet), *x^k^*.

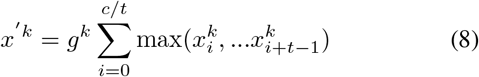

where c is the dimension of *x^k^*, and *t* is the 1D max pooling stride. In our experiments, *t* is chosen to be a power of 2 which sets the dimensions of 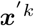 close to that of *e*. This is desirable as by setting the dimensions of the latent representations of each data modality to be close to the cross-modal representation, the network capacity for each distribution is normalized before being combined. The practical effect is that the complexity of relationships that are learnable for each data modality are made to be uniform.

Finally, the output of the fused network 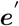 is the combination of both residual networks (Fig. 1, labeled part I: 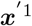 and part II: 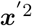), with the EmbraceNet output *e*. In our case, the terminal network is three dense layers of sizes 256, 128, and 2, sequentially.

### D. Performance metrics

#### 1) AUC-ROC score

To evaluate the performance of the proposed method for the seizure detection task, we used the area under the Receiver Operating Characteristic curve (AUC-ROC). The AUC-ROC measures the area under the recall vs. the false-positive rate (FPR) plots. Formal definitions of the recall and the FPR are provided below:

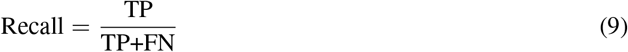

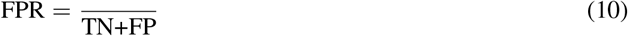

where TP, TN, FP, and FN represent true positives (correct seizure detection), true negatives (correct non-seizure detection), false positives (incorrect seizure detection), and false negatives (incorrect non-seizure detection).

#### 2) The Wilcoxon signed-rank test

To evaluate the performance of our model, the Wilcoxon signed-rank statistic is used. The obtained *p*-value provides a metric indicating the significance of performance improvement.

## IV. Results

### A. Test Cases

The following tests are applied to the TUH test set, with a pseudo-prospective study on the EPILEPSIAE and RPAH (selected and randomized) groups.

#### 1) Multimodal approach

To explore the effectiveness of our proposed network, we compare the following options, using the networks shown in Fig 1, on EEG and ECG recordings:

- ConvLSTM (EEG only)
- ResConv (ECG only)
- EmbraceNet (EN), prior state-of-the-art for multimodal data (EEG and ECG)
- Proposed (ECG and EEG)

#### 2) Missing modalities

The above networks are then tested for the cases where either EEG or ECG recordings are unavailable, which may occur due to poor electrode contact or unexpected signal dropout during recording.

##### a) Missing ECG (MC)

The ECG channel is treated as missing and is set to zero. The EEG channel is kept the same, and the same network models described above are used for testing (without retraining). Our proposed approach is compared to EmbraceNet, and the single-modal ConvLSTM EEG only network.

##### b) Missing EEG (ME)

All EEG inputs are set to zero while ECG recordings are used as per normal. The same comparison as the missing ECG case is made, but with the single-modal ResConv ECG only network instead.

### B. Performance

The distribution of AUC-ROC for the pseudo-prospective out-of-distribution analyses are shown in Fig. 2, tabulated in Table IV, and the ROCs under all test scenarios are provided in Fig. 5.

**Fig. 5:**
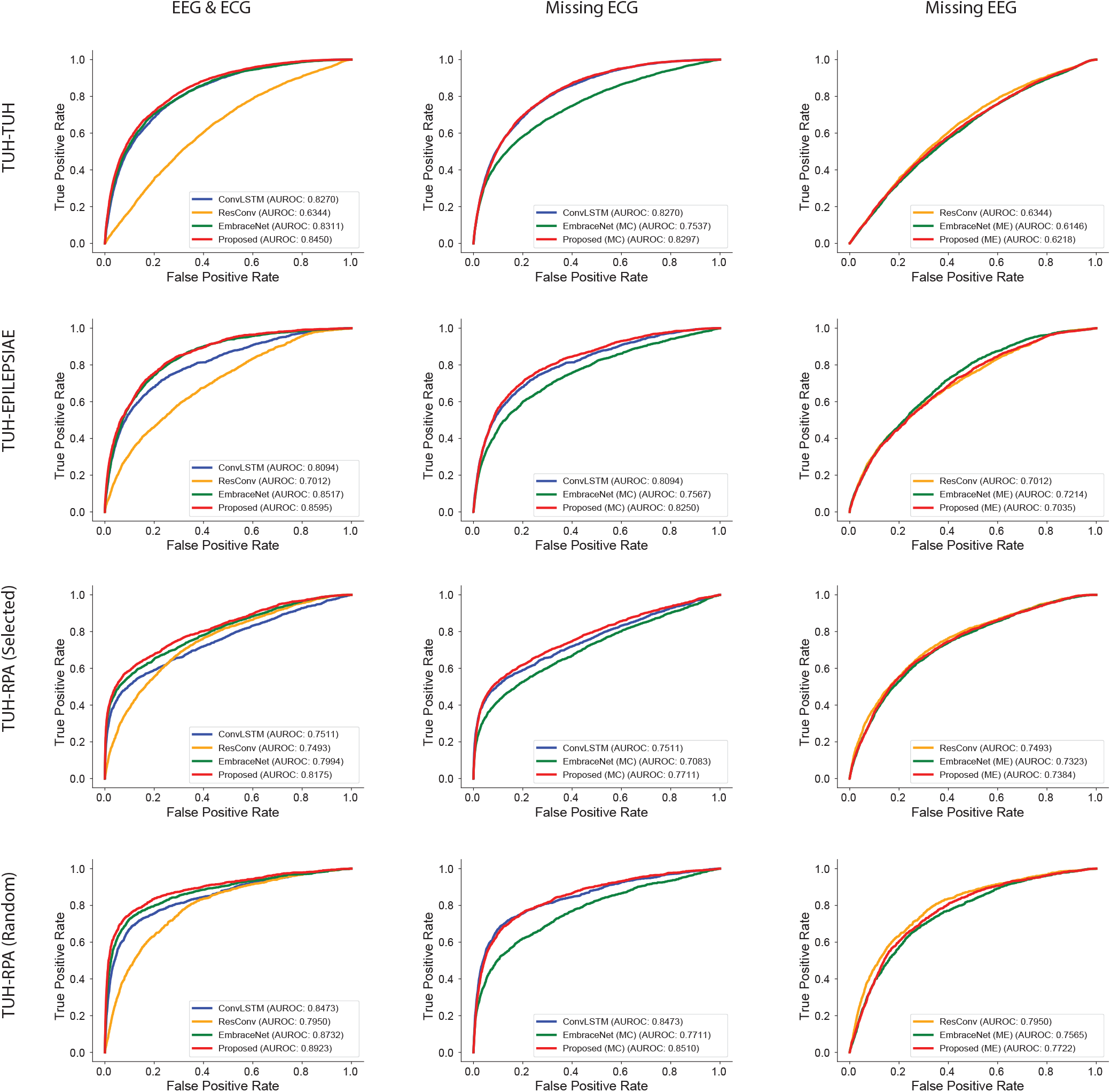
Receiver operating characteristic (ROC) curves for seizure detection. TUH-TUH, TUH-EPILEPSIAE, TUH-RPA (selected), TUH-RPA (random) use a model trained on the TUH training set, and assessed on the TUH test set, EPILEPSIAE dataset, RPA-selected and RPA-random sets. ConvLSTM: EEG-only with ConvLSTM network, ResConv: ECG-only with ResConv network, EmbraceNet: EmbraceNet architecture trained on combined EEG and ECG data, Proposed: our approach to combined EEG and ECG data, EmbraceNet (MC): EmbraceNet architecture trained on EEG and ECG data, but missing ECG during inference, EmbraceNet (ME): as above but missing EEG during inference, Proposed (MC): our proposed approach trained on EEG and ECG data, but missing ECG data during inference, Proposed (ME): as above, but missing EEG data during inference.

**TABLE IV:** Inference-only results comparison

| Inference-only test | Non-prospective | Pseudo-prospective |  |  |
| --- | --- | --- | --- | --- |
|  | TUH development | EPILEPSIAE | RPA-selected | RPA-random |
| EEG Hardware Manufacturer | Natus Medical Incorporated NicoletOne | Multiple devices | Compumedics Limited | Compumedics Limited |
| **Multimodal Input (Section IV-A1)** |  |  |  |  |
| ConvLSTM (EEG) | 0.8270 | 0.8094 | 0.7511 | 0.8473 |
| ResConv (ECG) | 0.6344 | 0.7012 | 0.7493 | 0.7950 |
| EmbraceNet (EEG+ECG) | 0.8311 | 0.8517 | 0.7994 | 0.8732 |
| Proposed (EEG+ECG) | **0.8450** | **0.8595** | **0.8175** | **0.8923** |
| **EEG-Only Input (Section IV-A2a)** |  |  |  |  |
| ConvLSTM (EEG) | 0.8270 | 0.8094 | 0.7511 | 0.8473 |
| EmbraceNet (MC) | 0.7537 (−8.86%) | 0.7567 (−6.51%) | 0.7083 (−5.70%) | 0.7711 (−8.99%) |
| Proposed (MC) | **0.8297** (+0.33%) | **0.8250** (+1.93%) | **0.7711** (+2.66%) | **0.8510** (+0.44%) |
| **ECG-Only Input (Sec. IV-A2b)** |  |  |  |  |
| ResConv (ECG) | **0.6344** | 0.7012 | **0.7493** | **0.7950** |
| EmbraceNet (ME) | 0.6146 (−3.12%) | **0.7214** (+2.88%) | 0.7323 (−2.27%) | 0.7565 (−4.84%) |
| Proposed (ME) | 0.6218 (−1.99%) | 0.7035 (+0.33%) | 0.7384 (−1.45%) | 0.7722 (−2.87%) |

EmbraceNet (MC): Missing ECG information to the EmbraceNet fused network, Proposed (MC): Missing ECG information to the proposed fused network, EmbraceNet (ME): Missing EEG information to the EmbraceNet fused network, Proposed (ME): Missing EEG information to the Proposed fused network

**TABLE V:** RPA-selected results

| Pat ID | ConvLSTM (EEG) | ResiConv (ECG) | EmbraceNet (EEG+ECG) | Proposed (EEG+ECG) | EmbraceNet (MC) | Proposed (MC) | EmbraceNet (ME) | Proposed (ME) |
| --- | --- | --- | --- | --- | --- | --- | --- | --- |
| 1 | 0.3168 | 0.9474 | 0.7667 | 0.8202 | 0.4113 | 0.5712 | 0.9086 | 0.9332 |
| 2 | 0.9145 | 0.8754 | 0.9236 | 0.9308 | 0.8650 | 0.9007 | 0.8316 | 0.8531 |
| 3 | 0.8446 | 0.8276 | 0.8794 | 0.9194 | 0.7633 | 0.8499 | 0.8248 | 0.7952 |
| 4 | 0.9649 | 0.7040 | 0.9507 | 0.9551 | 0.9126 | 0.8269 | 0.7247 | 0.6128 |
| 5 | 0.7499 | 0.7777 | 0.7924 | 0.8177 | 0.7035 | 0.7697 | 0.7114 | 0.7460 |
| 6 | 0.7068 | 0.7963 | 0.6872 | 0.8062 | 0.6829 | 0.7051 | 0.6937 | 0.6879 |
| 7 | 0.4558 | 0.8178 | 0.5868 | 0.6003 | 0.4502 | 0.5072 | 0.7543 | 0.7721 |
| 8 | 0.2626 | 0.5760 | 0.4695 | 0.3058 | 0.5504 | 0.1901 | 0.5969 | 0.5454 |
| 9 | 0.3330 | 0.4761 | 0.3527 | 0.4949 | 0.4913 | 0.4893 | 0.4365 | 0.4659 |
| 10 | 0.6079 | 0.6470 | 0.6800 | 0.7367 | 0.5131 | 0.6832 | 0.6516 | 0.6475 |
| 11 | 0.8888 | 0.8394 | 0.9092 | 0.9322 | 0.8186 | 0.9154 | 0.8261 | 0.8338 |
| 12 | 0.7970 | 0.5861 | 0.8439 | 0.8129 | 0.6921 | 0.7744 | 0.7404 | 0.6060 |
| 13 | 0.9372 | 0.8105 | 0.9668 | 0.9769 | 0.8815 | 0.9466 | 0.8290 | 0.8158 |
| 14 | 0.8349 | 0.9160 | 0.9346 | 0.9369 | 0.7083 | 0.8833 | 0.8846 | 0.9067 |
| 15 | 0.8774 | 0.8358 | 0.9450 | 0.9689 | 0.6927 | 0.9218 | 0.8296 | 0.8321 |
| 16 | 0.9682 | 0.6296 | 0.9659 | 0.9790 | 0.8799 | 0.9629 | 0.6373 | 0.6221 |
| 17 | 0.8069 | 0.6326 | 0.8533 | 0.8320 | 0.6952 | 0.7874 | 0.5132 | 0.5734 |
| 18 | 0.9711 | 0.8897 | 0.9853 | 0.9935 | 0.9017 | 0.9820 | 0.8797 | 0.8968 |
| 19 | 0.8273 | 0.8425 | 0.8942 | 0.9345 | 0.6893 | 0.9076 | 0.7979 | 0.8209 |
| 20 | 0.6143 | 0.5909 | 0.6475 | 0.6722 | 0.5649 | 0.6331 | 0.5639 | 0.5663 |
| 21 | 0.8964 | 0.7536 | 0.9140 | 0.9156 | 0.8496 | 0.9056 | 0.7172 | 0.7210 |
| 22 | 0.9182 | 0.9284 | 0.9826 | 0.9815 | 0.8119 | 0.9596 | 0.9310 | 0.9334 |
| 23 | 0.6681 | 0.7673 | 0.6955 | 0.7346 | 0.6067 | 0.6985 | 0.6868 | 0.7322 |
| 24 | 0.7694 | 0.5914 | 0.7508 | 0.8236 | 0.6624 | 0.7815 | 0.6143 | 0.6069 |
| 25 | 0.6463 | 0.4647 | 0.6787 | 0.7018 | 0.6764 | 0.7133 | 0.5195 | 0.4715 |
| 26 | 0.7256 | 0.7661 | 0.7703 | 0.7822 | 0.6681 | 0.6953 | 0.7098 | 0.7346 |
| 27 | 0.5520 | 0.6834 | 0.6751 | 0.6433 | 0.5848 | 0.5938 | 0.6734 | 0.6853 |
| 28 | 0.7250 | 0.8971 | 0.8694 | 0.8901 | 0.7534 | 0.7790 | 0.8835 | 0.8955 |
| 29 | 0.3986 | 0.7851 | 0.6252 | 0.6417 | 0.4710 | 0.5041 | 0.8243 | 0.7890 |
| 30 | 0.9203 | 0.6220 | 0.8949 | 0.8921 | 0.8997 | 0.8863 | 0.4956 | 0.5191 |
| 31 | 0.3942 | 0.3756 | 0.3623 | 0.3678 | 0.5205 | 0.4929 | 0.3731 | 0.3637 |
| Overall | 0.7511 | 0.7493 | 0.7994 | 0.8175 | 0.7083 | 0.7711 | 0.7323 | 0.7384 |

**TABLE VI:** EPILEPSIAE dataset results

| Pat ID | ConvLSTM (EEG) | ResConv (ECG) | EmbraceNet (EEG+ECG) | Proposed (EEG+ECG) | EmbraceNet (MC) | Proposed (MC) | EmbraceNet (ME) | Proposed (ME) |
| --- | --- | --- | --- | --- | --- | --- | --- | --- |
| 1 | 0.8843 | 0.6913 | 0.8967 | 0.926 | 0.7752 | 0.9200 | 0.7055 | 0.6942 |
| 2 | 0.8876 | 0.4561 | 0.9200 | 0.9058 | 0.8455 | 0.8869 | 0.5677 | 0.5115 |
| 3 | 0.8596 | 0.8838 | 0.9292 | 0.9293 | 0.7740 | 0.8810 | 0.9135 | 0.9074 |
| 4 | 0.9311 | 0.5172 | 0.9153 | 0.9149 | 0.8736 | 0.9274 | 0.5680 | 0.5283 |
| 5 | 0.7449 | 0.5888 | 0.8159 | 0.8023 | 0.7542 | 0.7442 | 0.6331 | 0.6142 |
| 6 | 0.7 | 0.7311 | 0.7163 | 0.7402 | 0.6570 | 0.6534 | 0.6639 | 0.6858 |
| 7 | 0.8352 | 0.785 | 0.8801 | 0.893 | 0.6873 | 0.7982 | 0.8244 | 0.7811 |
| 8 | 0.8266 | 0.9023 | 0.9131 | 0.8979 | 0.7876 | 0.8844 | 0.8851 | 0.8736 |
| 9 | 0.9347 | 0.6483 | 0.9245 | 0.9395 | 0.9182 | 0.9450 | 0.6504 | 0.6387 |
| 10 | 0.6768 | 0.6737 | 0.8251 | 0.8094 | 0.7171 | 0.7467 | 0.7274 | 0.7000 |
| 11 | 0.8379 | 0.73 | 0.8595 | 0.8879 | 0.7220 | 0.8794 | 0.7239 | 0.7047 |
| 12 | 0.8476 | 0.795 | 0.9321 | 0.9332 | 0.7830 | 0.8670 | 0.7940 | 0.8001 |
| 13 | 0.9009 | 0.7533 | 0.9114 | 0.9281 | 0.8371 | 0.9124 | 0.7873 | 0.7652 |
| 14 | 0.8839 | 0.7015 | 0.9271 | 0.9183 | 0.8234 | 0.8860 | 0.7503 | 0.7192 |
| 15 | 0.708 | 0.7452 | 0.7650 | 0.774 | 0.7345 | 0.7017 | 0.7318 | 0.7119 |
| 16 | 0.8931 | 0.7481 | 0.8974 | 0.9021 | 0.7757 | 0.8480 | 0.7615 | 0.7182 |
| 17 | 0.8322 | 0.7481 | 0.9004 | 0.8938 | 0.7354 | 0.8351 | 0.7776 | 0.7534 |
| 18 | 0.7605 | 0.846 | 0.8476 | 0.8663 | 0.6995 | 0.7495 | 0.8373 | 0.8535 |
| 19 | 0.929 | 0.7892 | 0.9295 | 0.9415 | 0.8359 | 0.9147 | 0.8441 | 0.7994 |
| 20 | 0.6692 | 0.8621 | 0.8604 | 0.8581 | 0.6746 | 0.7141 | 0.8922 | 0.8701 |
| 21 | 0.8692 | 0.7291 | 0.8955 | 0.9047 | 0.8282 | 0.9044 | 0.7961 | 0.7472 |
| 22 | 0.7439 | 0.761 | 0.8479 | 0.8338 | 0.7513 | 0.7630 | 0.7818 | 0.7646 |
| 23 | 0.9715 | 0.7369 | 0.9735 | 0.9801 | 0.8911 | 0.9670 | 0.7873 | 0.7533 |
| 24 | 0.7528 | 0.6739 | 0.7526 | 0.7843 | 0.6545 | 0.7515 | 0.7411 | 0.7161 |
| 25 | 0.7815 | 0.6334 | 0.8554 | 0.8409 | 0.7665 | 0.7871 | 0.6456 | 0.6290 |
| 26 | 0.6157 | 0.5805 | 0.6227 | 0.675 | 0.6110 | 0.6743 | 0.5828 | 0.5726 |
| 27 | 0.8838 | 0.8691 | 0.9390 | 0.9381 | 0.7682 | 0.8690 | 0.8758 | 0.8609 |
| 28 | 0.905 | 0.7212 | 0.8809 | 0.8893 | 0.8104 | 0.8741 | 0.7045 | 0.7075 |
| 29 | 0.6815 | 0.7006 | 0.8110 | 0.829 | 0.6882 | 0.7939 | 0.7251 | 0.7213 |
| 30 | 0.7842 | 0.6461 | 0.8485 | 0.8472 | 0.7053 | 0.8485 | 0.6859 | 0.6316 |
| Overall | 0.8094 | 0.7012 | 0.8517 | 0.8595 | 0.7567 | 0.825 | 0.7214 | 0.7035 |

**TABLE VII:**
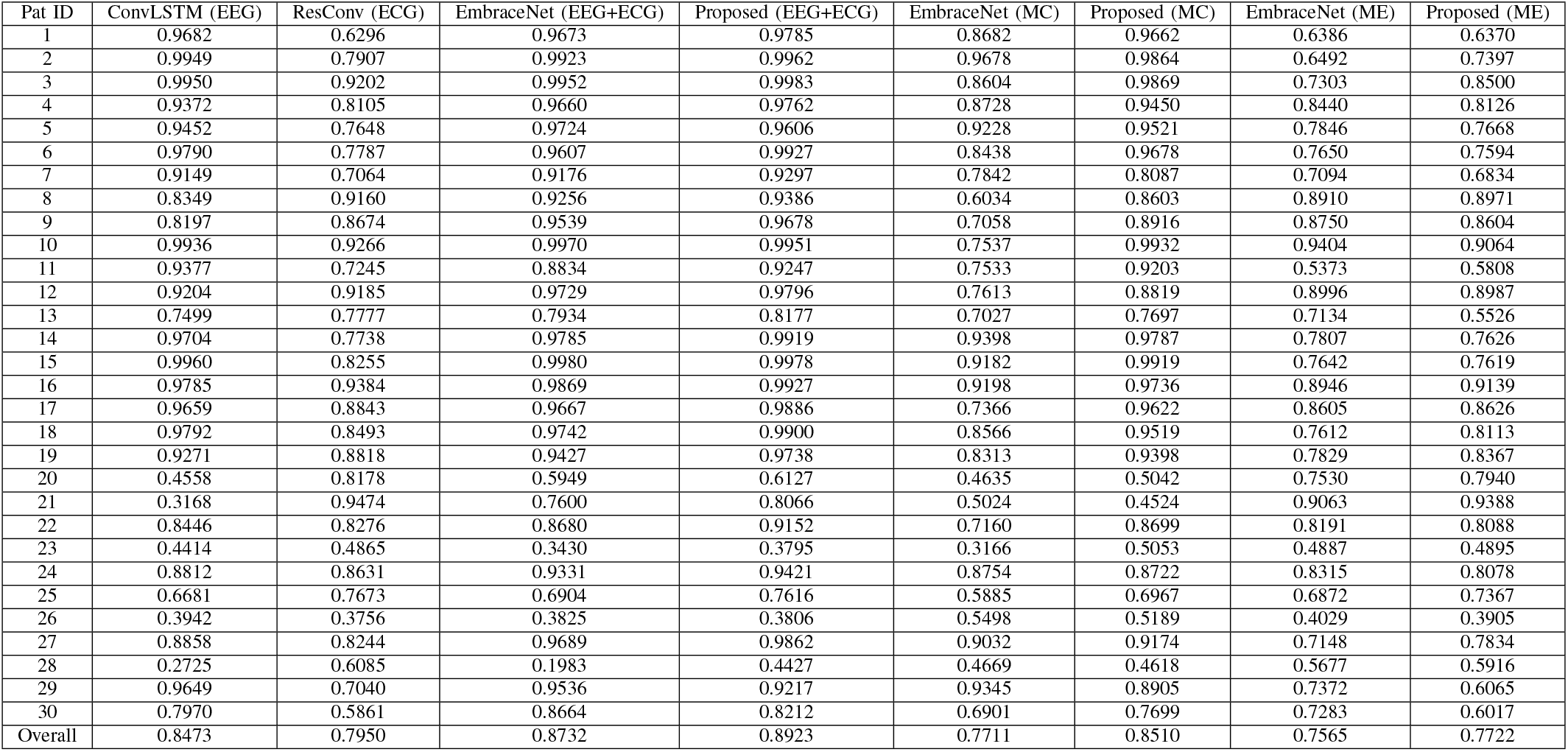
RPA-random results

The AUC-ROC for each individual patient across the EPILEPSIAE, RPA-selected and randomized groups is shown in Fig. 2. The first column depicts the results for the multimodal approach. Combining EEG and ECG data using a fused network (for both EmbraceNet and our proposed method) improves performance across all three out-of-distribution datasets. Our proposed method consistently outperforms EmbraceNet across the three experiments. From Table IV, the absolute AUC improvement of our proposed method when compared with EmbraceNet is 0.014 for the TUH test set, 0.008 for the ELIEPSIAE set, 0.018 for the selected RPA group, and 0.019 for the randomized RPA group.

The absolute margin of the AUC-ROC from our approach improves upon the prior state-of-the-art on the out-of-distribution datasets by 6.71% for EEG-only, 14.42% for ECG-only, and 1.76% for EmbraceNet’s multimodal approach. The AUC-ROC curves across each dataset are shown in Fig. 5. The *p*-value derived from the Wilcoxon signed-rank test for our proposed model (EEG+ECG) when compared with EEG only, ECG only and EmbraceNet (EEG+ECG) is (*p* < 0.0001, *p* < 0.0001, *p* = 0.04182) in the EPILEPSIAE dataset, (*p* < 0.0001, *p* = 0.00368, *p* = 0.00126) for the selected RPA group, and (*p* = 0.00011, *p* = 0.00056, *p* = 0.00028) RPA-random group. This demonstrates the statistically significant performance of our model for a specified threshold of 0.01.

For the test case of the missing ECG modality (MC), we repeat the above tests for when the two fused networks (our proposed and EmbraceNet) are missing ECG information. Compared with EEG-only performance using the ConvLSTM network, the performance of EmbraceNet dropped by 8.86% (TUH), 6.51% (EPILEPSIAE), 5.70% (RPA-selected), and 8.99% (RPA-random). In contrast, our proposed method increases performance of the ConvLSTM EEG-only network by 0.33% (TUH), 1.93% (EPILEPSIAE), 2.66% (RPA-selected), and 0.44% (RPA-random). The *p*-value under this case for our proposed approach against EmbraceNet is (*p* < 0.0001, *p* = 0.00013, *p* < 0.0001 EPILEPSIAE, RPA-selected and -random datasets, respectively.

The above process was repeated for the missing EEG modality case (ME). When compared with the ECG-only Resi-Conv network, the performance of EmbraceNet with missing EEG information dropped by 3.12% (TUH), 2.27% (RPA-selected), and 4.84% (RPA-random) and improved by 2.88% (EPILEPSIAE). The proposed method dropped by 1.99% (TUH), 1.45% (RPA-selected), and 2.87% (RPA-random), and improved by 0.33% (EPILEPSIAE). The *p*-values when compared to EmbraceNet (ME) *p* = 0.26435 (RPA-selected), and *p* = 0.38974. (RPA-random). Other than the EPILEPSIAE dataset, the decrease in our model’s performance is less than the decrease in EmbraceNet’s performance, demonstrating it is robust when the EEG modality is missing.

## V. Discussion

For the multimodal case, described in Section IV-A1, our proposed model outperforms the AUC-ROC for seizure detection compared to the EEG-only network, ECG-only network, and the EmbraceNet fused approach across all datasets.

Our approach on the non-prospective TUH test set improved the AUC-ROC attained using EmbraceNet from 0.8311 to 0.8450. This improvement of 1.67% is the smallest margin of all cases. All pseudo-prospective trials showed a larger improvement other than for EPILEPSIAE (0.008), thus demon-strating our method’s capacity for generalization and potential for use in a clinical setting. The ECG-only network was the only approach to improve upon the TUH test set baseline on pseudo-prospective samples, but came with the cost of large performance variation and significantly lower absolute AUC-ROC. We expect this variation arises from individuals with different seizure origins which variably influence the autonomic nervous system [45]. The table shows that performance on the RPA-random set is better than the RPA-selected by 9.15% when using the proposed method to test. This may be due to most patients’ seizure foci are on the temporal lobe in the RPA dataset, and patients with seizure foci on the temporal lobe typically show higher performance than other locations.

For the ECG-missing case, described in Section IV-A2a, we evaluated the AUC performance when all ECG data was omitted. Our results show that the proposed method is still able to marginally improve the performance over the EEG-only network (average improvement of 1.30%), whereas the EmbraceNet approach has a significant drop in performance (average drop of 7.51%). This reflects that our proposed network has high robustness for missing (potentially corrupted) ECG recordings.

Finally, we analyze the case for missing EEG information, discussed in Section IV-A2b. Our proposed method experienced a slight drop in performance when compared to the dedicated ECG-only ResConv model on the TUH and RPAH groups, and an increase on the EPILEPSIAE dataset by 0.33%. The performance of EmbraceNet dropped more when compared (in the range of 2.27% and 4.84%) other than for the EPILEPSIAE dataset. Our network is much more stable than EmbraceNet, although this test makes it evident that our network relies more on EEG data to surpass the other networks.

Our results illustrate that our proposed model and method are generalizable across different countries’ datasets which use different recording equipment. We have also shown the robustness of our network in terms of performance and susceptibility to missing data modalities when compared to the state-of-the-art in data fusion. Our experiments also prove that using multimodal data can achieve better performance than either EEG or ECG alone.

## VI. Conclusion

Despite the extensive studies over the past four decades of using EEG in seizure detection, the use of ECGs is quite limited and never previously reported in a multimodal deep learning model. our proposed model and fused modality approach show promise in using EEG and ECG signals together, and were demonstrated to generalize to pseudo-prospective studies. our analysis shows that a seizure detection system can sustain state-of-the-art performance on out-of-distribution samples, which is a critical feature for clinical translation.

## Acknowledgements

Yikai Yang would like to acknowledge the Research Training Program (RTP) support provided by the Australia Government. Omid Kavehei acknowledges the support provided by The University of Sydney through a SOAR Fellowship and Microsoft’s support through a Microsoft AI for Accessibility grant.

## Ethics declarations

Ethics *X*19-0323-2019/STE16040: Validating epileptic seizure detection, prediction and classification algorithms approved on 19 September 2019 by the NSW Local Health District (LHD), for implementation in the Comprehensive Epilepsy Services, Department of Neurology, The Royal Prince Alfred Hospital.

## Data availability

The Temple University Hospital dataset is publicly available at https://www.isip.piconepress.com/projects/tuh_eeg/html/downloads.shtml. The training on the publicly available TUH dataset provides readers with sufficient level of ability to independently confirm some of the results reported. The EPILEPSIAE dataset is available at cost via http://www.epilepsiae.eu/projeck_outputs/european_database_on_epilepsy. The Royal Prince Alfred Hospital was used under ethics Review Board approval for our use only.

## Code availability

The code used to generate all results in this manuscript can be made available upon request.

## Supplementary information

